# Computational Analysis of Microtubule-Mediated Saltatory Neuroelectrical Transmission

**DOI:** 10.64898/2026.02.25.708076

**Authors:** Yong Xiao Yang, Bao Ting Zhu

## Abstract

It was recently postulated that neural microtubules (neuro-MTs), which are densely packed inside axons and dendrites, are vacuum cylindrical nanotubes that can mediate neuroelectrical transmission with a unique form of quasi-superconductivity. In this work, the behaviors of free electrons inside a theoretical neuro-MT are modeled using computational analysis and calculations. We reveal that neuro-MTs can function as nanosized physiological devices that mediate neuroelectrical transmission with a super-high energy efficiency. Under physiological conditions, the binding of cytosolic cations (*e*.*g*., K^+^ and Na^+^) to the surface residues of a neuro-MT triggers its transition from the resting state to an active state, and the rapid dissociation of these cations triggers the opposite. The dipole ring structures of a neuro-MT will help terminate the free electron conduction inside with high efficiency. The proposed neuro-MT-mediated electrical transmission offers a novel mechanistic explanation for the saltatory conduction of the action potentials along an axon. This study also provides insights into the design of novel biomimicking room-temperature superconducting materials, such as the quasi-superconducting carbon or silicone nanotubes.

## INTRODUCTION

The nervous system, brain in particular, conducts massive amount of neuroelectrical transmission almost all the time, even during sleep. Unlike man-made electronic devices such as computers, the nervous system surprisingly almost never suffers from “overheating” due to its overwhelming amount of neuroelectrical transmissions. It was postulated in recent years that neuroelectrical transmission in the nervous system likely is mediated by a mechanism of quasi-superconductivity which takes place under physiological temperatures and conditions (1).

At present, the Bardeen–Cooper–Schrieffer theory is widely accepted to explain the mechanism of superconductivity of regular metallic or composite superconductors, especially under low temperatures (2). Recently, a new hypothesis concerning the physical properties and requirements of superconducting materials was proposed (3). It was postulated that for superconductivity (*i*.*e*., with zero electrical resistivity) to occur, the conductor must have nanosized straight vacuum tunnels inside with radius size large enough to allow the passage of free conduction electrons (likely in a ballistic manner) without collisions. In addition, some of the composite atoms of the conductor should be able to readily release free electrons to form the conduction band. This hypothesis is supported by some experimental observations in the literature, and also offers a plausible explanation why carbon nanotubes or graphene sheets can become superconducting (or quasi-superconducting) under certain conditions as these nano-devices contain ample vacuum spaces inside to enable mostly collision-free passage of the conduction electrons (discussed in detail in ref. [1]).

The above hypothesis on superconductivity (3) has also led to the speculation that the neuro-MTs, which are major structural components of axons and dendrites, may function as unique nanosized biodevices that can mediate electrical transmission with a quasi-superconducting property (1). Structurally, MTs are linear cylindrical tubes made of tubulin polymers (4), consisting of α- and β-tubulin subunits (4-6). Structurally, each MT usually consists of 13–15 rows of protofilaments (7) made of α-/β-tubulin heterodimers, and the protofilaments are associated laterally and closed into a nanotube (8-10). Inside the axons and dendrites, there are large numbers of densely-packed MTs, forming a distinct structure. It was hypothesized in 2022 that the MTs are involved in neuroelectrical transmission, and these MTs are referred to as “neuro-MTs” (1). Structurally, it was hypothesized that neuro-MTs have a hollow and vacuum structure inside which facilitates the collision-free slow passage of the free conduction electrons (1).

In the past thirty years, some experimental structures of MTs or proteins related to MTs have been determined using X-ray crystallography or cryo-EM methods (5,11-13). There were also studies exploring their electrostatic or electromagnetic properties (1, 14-42). In the present study, computational analysis is adopted to model the electric fields (EFs) and the behavior of free electrons inside a seamless neuro-MT, and based on the knowledge gained from the computational modeling analysis, a new mechanistic explanation on the saltatory neuroelectrical transmission is tendered.

## MATERIALS AND METHODS

### Experimental structure of a seamless MT

The experimental structure of a seamless MT was determined earlier using cryo-EM by Benoit *et al*. (11). The structural file of MT (PDB code: 6b0i, CIF file: 6b0i-assembly1.cif) was downloaded from the Protein Data Bank (http://www.rcsb.org/) (43). The missing residues in the structures of tubulin α (residues 442 □ 451) and β (residues 430 □445) were added based on the AlphaFold predicted structures (44).

### Estimation of the electric field direction and its relative intensity

In order to estimate the electric field (EF) direction and relative intensity inside a theoretical MT tunnel, the atomic charges in the MT structure are assigned based on the *CHARMM36m* force field (45) using *CHARMM-GUI* (http://www.charmm-gui.org) (46). The length of a theoretical MT was expanded to a micrometer scale by repeating the coordinates of the known structures along the central axis. For ease of calculation, each amino acid residue, GDP or GTP in the MT structure is individually represented using the geometric center of the corresponding molecule, and the charge is the sum of all atomic charges. The EF direction and relative intensity are estimated according to the Coulomb electrostatic interactions. The relative intensity and direction of the summated EF at a given point inside the MT tunnel is calculated using the following equation:

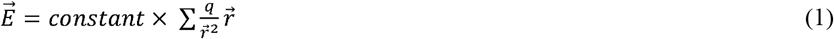

Here, 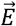 and *q* are the relative EF intensity and the charge of a residue, respectively. 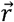 is the vector from the geometric center of a residue to the given point. ∑ represents the sum traversing all the charged residues in an MT, plus all GDP (–2), GTP (–3) and Mg^2+^ (+2).

### Simulating the movement of a free electron inside a neuro-MT

The force exerted on a free electron inside a neuro-MT is computed using the following formula:

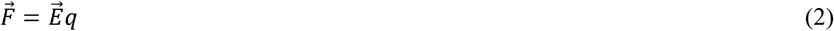

Here, 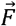, *q* and 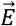 refer to the force, mass of the charged particle (*i*.*e*., electron), and the EF intensity, respectively. When a free electron moves along inside the MT tunnel, its relative velocity is calculated according to Newton’s second law, which is expressed as follows:

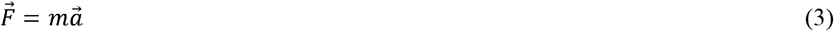

Here, 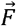, *m* and 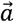 are the force, mass and accelerated velocity of the charged particle, respectively.

When the displacement distance 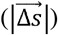 and time (Δ*t*) are very small, the instantaneous accelerated velocity can be regarded as a constant and represented using the velocity at the initial position. Assuming that the charged particle moves from the initial position 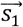 to position 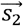, the time (Δ*t*), the instantaneous speeds at the two positions (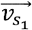and 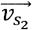), theacceleration velocity 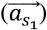, and the displacement vector 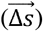 (from the positions 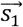to 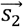) satisfy the following quantitative relationships:

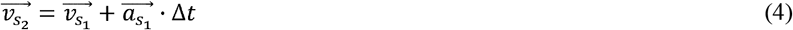

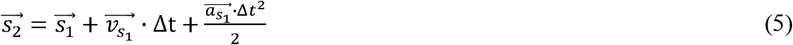

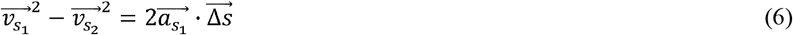

Here, the direction of the instantaneous speeds (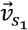 and 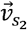), the acceleration velocity 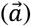and the displacement vector 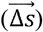are on the same straight line. The instantaneous positions along the trajectory of the kinetic mass point are derived from the above equations.

## RESULTS AND DISCUSSION

### Overall structure and charge distribution of a neuro-MT

Presently, the complete structure of a neuro-MT is still unclear. The 15-membered seamless MT (11) is adopted in this study as a prototype neuro-MT for modeling analysis of its biophysical properties; its structure is shown in **Suppl. Fig. S1A** (side view) and **Suppl. Fig. S1B** (top view). Two chains of the αβ dimers intertwine each other which are somewhat similar to the two chains in the double helical DNA structure (**Suppl. Fig. S1A**). The vertical length of two αβ dimers is approximately 165 □ (**Suppl. Fig. S1A**), and the inside diameter of the tunnel inside a 15-membered MT is approximately 95 Å (**Suppl. Fig. S1B**).

According to the assigned charges, there are 103 charged amino acid residues (41 positive charges, 62 negative charges), 1 GTP (–3) and 1 Mg^2+^ (+2) in tubulin α; there are 99 charged residues (38 positive charges, 61 negative charges) and 1 GDP (–2) in tubulin β. Accordingly, the total net charges are –22 in tubulin α and –25 in tubulin β. The overall charge distributions are shown in **Suppl. Fig. S1C** (side view) and **Suppl. Fig. S1D** (top view). The overall distribution of the positive and negative charges is very similar for both tubulin α and β helical structures (**Suppl. Fig. S2A, S2B**). Notably, both tubulin α and β subunits contain markedly more negative-charged amino acids than positive-charged amino acids, and particularly, the outermost part of a neuro-MT contains mostly negative-charged amino acids (**Suppl. Fig. S1D, Suppl. Fig. S2C, S2D**). This unique feature enables a neuro-MT to attract and bind intracellular cations (such as K^+^ and Na^+^) and thus become a reservoir (“sink”) for these cations.

### A theoretical neuro-MT at the resting state

It was recently hypothesized that a neuro-MT will allow the conduction of free electrons upon nerve stimulation (1). In this study, first we simulated the EF direction and relative intensity inside a neuro-MT according to Eq. 1. During the initial modeling analysis, the length of a neuro-MT is expanded to the micrometer scale by repeating the MT period which contains 30 tubulin α and β subunits (as illustrated in **Suppl. Fig. S1A**) 100 times.

To analyze the effect of the MT’s charge distribution on the EF inside, the charge of the residues at the outer and inner regions of a neuro-MT is manually altered. Here, the outer surface region refers to the region with a radius >140 Å; the inner surface region refers to the region with a radius <110 Å; and the middle region refers to the region with a radius between 110 and 140 Å (depicted in **Suppl. Fig. S1D**).

Under normal physiological condition when a neuro-MT is at the resting state, it is believed that the EF intensities inside its tunnel theoretically should be zero along the central axis such that all free electrons inside the tunnel will be static, likely attaching to the inner surface wall (explanation provided later). To model the EF intensities inside of a neuro-MT at the resting state, free electrons are placed at the inner surface region, and the net charge of all positive-charged positions at the inner surface region will become zero (*i*.*e*., they are neutralized by the free electrons present inside a neuro-MT). As for the sources of free electrons present inside a neuro-MT, it was postulated earlier that they are formed by tubulins using GTP molecules as the energy source (1). Additionally, it should be noted that each negative-charged position at the outer surface region of a neuro-MT will be altered according to the change in the intraneuronal cation concentrations during an action potential (AP).

As shown in **Fig. 1A**, a series of initial calculations were performed when each negative-charged position on the outer surface of a neuro-MT was assigned to carry a certain amount of charge (ranging from +0.16 to +0.21). We found that the EF intensities inside a neuro-MT’s tunnel will become near-zero along the central axis when each negative-charged position on the outer surface carries an average charge of +0.17 (**Fig. 1A**). Notably, +0.17 average charge would mean that roughly 17% of all negative-charged amino acid residues on the outer surface carry a net charge of +1 (such as carrying a K^+^ ion) while all other negative charged residues are neutralized by a cation such as K^+^ (thus carrying zero charge). This condition is highly feasible and relevant physiologically. Indeed, after all negative-charged residues on the outer surface are neutralized and then an additional 17.0% (240/1412) of the residues at the outmost region of a neuro-MT are bound with a cation (such as K^+^ ion) and carry a net charge of +1, the EF intensities along the central axis become literally zero (**Fig. 1B, 1C**). This modeling result reveals that under certain physiologically-relevant conditions, the EF inside a neuro-MT will disappear along the central axis, which means that the free electrons inside a neuro-MT will be static (not moving in either direction). Notably, under this “resting state” condition, the modeling results show that there is still a small EF projected toward the central axis on the cross sections (*i*.*e*., the ***x–y*** planes) (**Fig. 1D**). This small EF on the cross section will force all free electrons (which carry a negative charge) to move toward the inner surface region, where they would be in close association with the positive-charged inner surface residues due to Columbus interactions.

**Figure 1.**
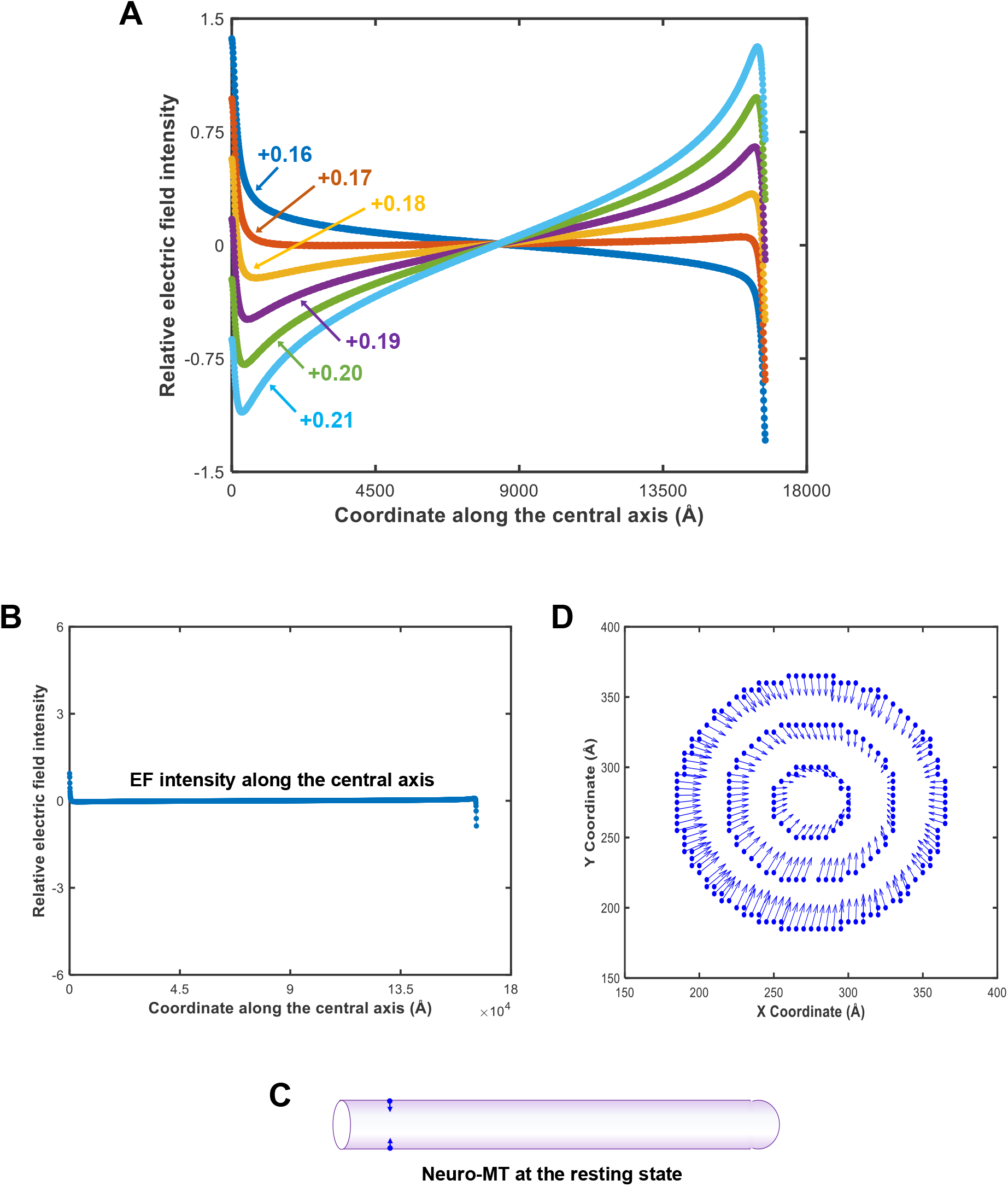
Modeling of the electric fields (EFs) inside a neuro-MT at the resting state. **A**. Modeling of the relative magnitude of EFs on the central axis (*z*-axis) inside a neuro-MT. During the modeling analysis, each of the outside negative-charged amino acid residues is assigned to carry a certain amount of average charge ranging from +0.16 to + 0.21), and then the relative magnitude and directions of the EF on the central axis are calculated. **B**. Relative EF intensity on the central axis of a neuro-MT at the resting state where the negative-charged residues on the outer surface are neutralized and 17.0% (240/1412) of these residues are bound with a K^+^ ion (*i*.*e*., carrying a net charge of +1). **C**. EF directions and relative intensities inside a neuro-MT at the resting state. The blue arrows depict the EF directions at relevant positions, and the arrow length represents the relative magnitude of the EF intensity. **D**. EF directions and relative intensities (based on arrow length) on the cross-section at the resting state.

### Electron movements inside a theoretical neuro-MT at the active state

During neuroelectrical transmission, it is known that only a small segment of a myelinated axon (or dendrite), which is called “the node of Ranvier (RN)”, will experience rapid influx of cations (mostly Na^+^ from the extracellular space) when an AP arrives at the NR. As such, only a small segment in the long neuro-MT will have additional cations placed at the outer surface residues during neurotransmission. Next, we choose to model the change in EF intensity and direction after we alter the outer surface charge in a small fraction of a neuro-MT by letting about 36.4% (514/1412) of the outer surface residues carry a net charge of +1 while the remainder residues neutralized. For this modeling analysis, the length of the neuro-MT is enlarged by repeating the earlier segment (which produced the modeling data in **Fig. 1B–1D**) 10 times again. The mechanism of electrotransmission inside a neuro-MT is deduced by analyzing the EF and the conduction of free electrons inside a neuro-MT’s tunnel.

Based on computational modeling, there is an EF generated along the central axis during the active state (**Fig. 2A**), and the EF points in the opposite directions on the central axis away from the “clamped” segment (depicted in **Fig. 2B**). This means that the free electrons inside a neuro-MT will move toward the clamped segment (*i*.*e*., the segment which has cations placed on its outer surface). Additionally, there is an EF on the cross sections (*i*.*e*., the ***x–y*** planes) of the neuro-MT, and its directions are projected away from the central axis (**Fig. 2C**). This EF on the ***x–y*** planes will force the free electrons to leave the inner surface region and move toward the center region of the tunnel. Note that the above modeling analysis was also performed when different fractions of the negative-charged residues carry a +1 positive charge (*i*.*e*., bound by a Na^+^ ion). We found that essentially the same observations were made when varying fractions of the negative-charged residues are carrying a positive charge.

**Figure 2.**
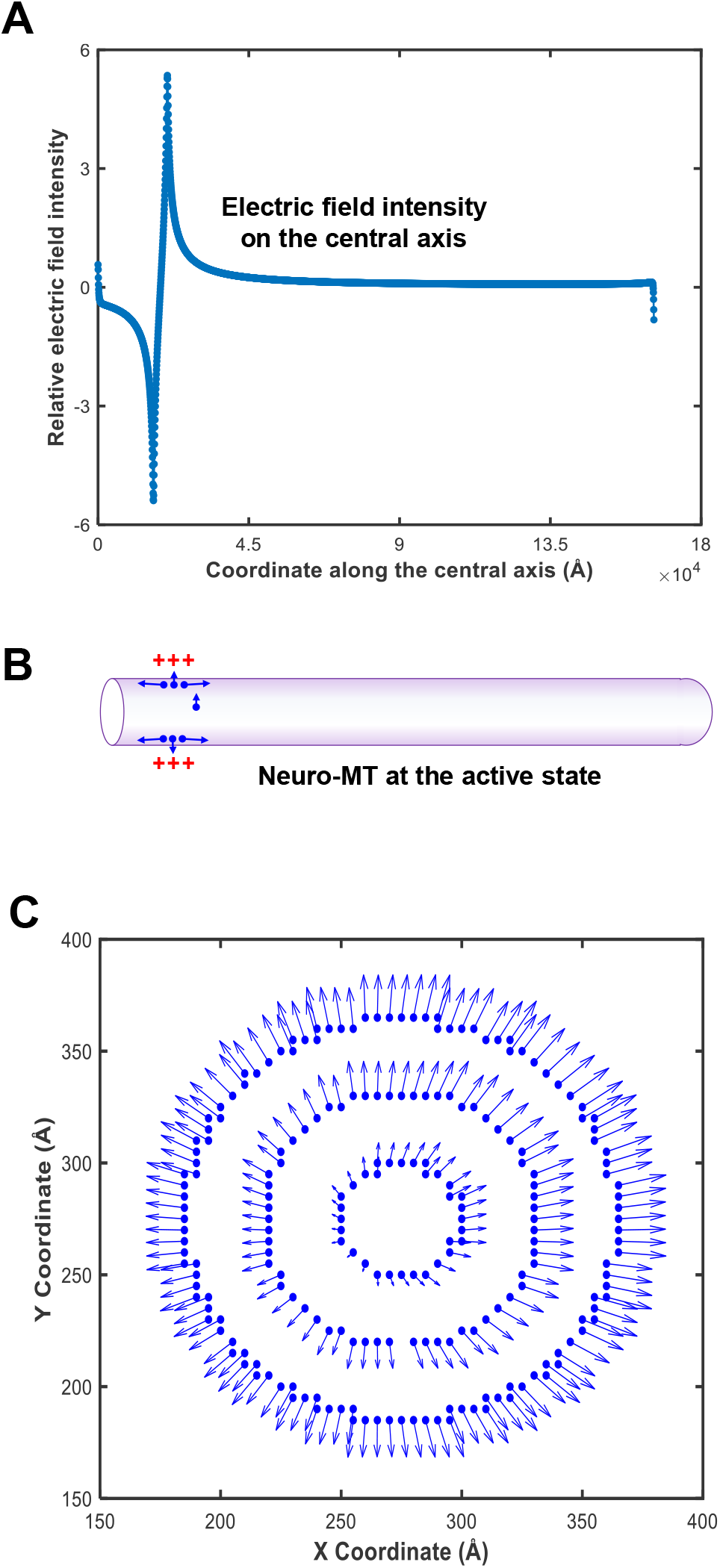
Modeling of the electric fields (EFs) inside a neuro-MT at the active state. **A**. Relative EF intensities on the central axis of a neuro-MT during the active state. To mimic the active state, approximately 36% of the outer surface residues in a small fraction of a neuro-MT (*i*.*e*., the NR region) are allowed to carry a net charge of +1 while other remainder residues are neutralized (*i*.*e*., without carrying any charge). **B**. Schematic depiction of the EF directions and relative intensities inside a neuro-MT during the active state. The blue arrows depict the EF directions at relevant positions, and the arrow length represents the relative magnitude of the EF intensity. **C**. EF directions and relative intensities (based on arrow length) on the cross-section at the active state.

Next, we model the relative movements of 6 representative electrons with different initial positions along the central axis of a neuro-MT (depicted in **Fig. 3A**) when it is changed from the resting state to the active state, when an AP is generated at node of Ranvier 1 (NR1), and then passes along from NR1 to NR2 (**Fig. 3A**). Here, we assume that all 6 electrons are static in their initial positions during the initial resting state (i.e., before the AP is generated at NR1). The velocity and relative positions of the free conduction electrons along the central axis are estimated according to Eq. 4–6.

**Figure 3.**
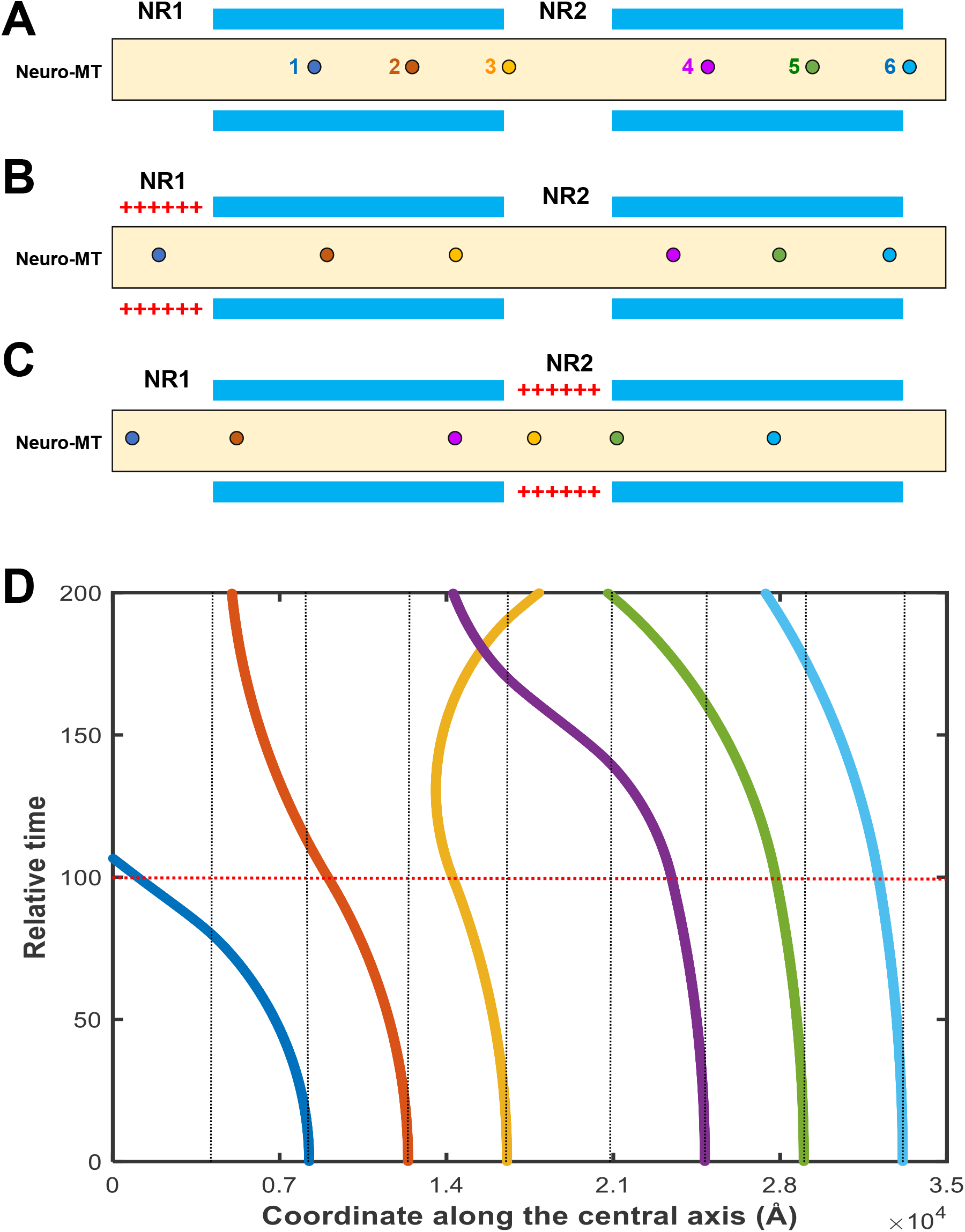
Modeling of the movement of free electrons inside the vacuum tunnel of a neuro-MT. **A**. Schematic depiction of the six free electrons inside the vacumn tunnel of a neuro-MT at the initial resting state. The blue colored bars represent the regions where the myelin sheath is located. Two nodes of Ranvier (NR1 and NR2) are drawn. At the resting state, the EF is zero inside the neuro-MT tunnel, and the electrons are static. **B**. When an action potential (AP) is generated at NR1, the clamping voltage will have a drawing force on all six free electrons in the same direction. The closer of the electron to the clamping voltage, the stronger the drawing force. **C**. When the AP moves to NR2, the clamping voltage at NR1 is expected to disappear very quickly. The forces generated by the “clamping voltage” at NR2 will exert a drawing force on the free electrons on both sides of NR2 in opposite directions. **D**. Quantitative analysis of the positions of the six free electrons inside the tunnel of a neuro-MT during the initial 200 time units (note that an AP will stay in a NR for only 100 time units and will then move to the next NR). Here, it should be noted that when the AP jumps to the next NRs, the dispperance of the clamping voltage at the preceding NR may quickly stop moving in either directions due to the EFs there which will force the free elections to move toward the inner surface and stay there. In the modeling results shown here, all the free electrons at the NR1 are allowed to continue to move according to the forces acting on them when the AP jumps to NR2.

As depicted in **Fig. 3A**, when cations (mostly Na^+^ and K^+^) are placed at the outer surface of a small fraction of a neuro-MT (such as NR1), it is just like placing a “clamping voltage” on this region of a neuro-MT, mimicking the active state when an AP is generated at NR1. Understandably, the forces acting on the free electrons closer to the positive-charged region are bigger than the forces acting on the electrons farther away from the charged region. Therefore, when NR1 is activated and before it jumps to NR2 (here the relative action time is set to 100 for each NR activation) (**Fig. 3B, 3D**), electron 1 moves toward NR1 fastest, electrons 2 and 3 move toward NR1 at reduced speeds, and electrons 4–6 may barely move (due to the weaker forces acting on them). When the activation reaches NR2 (assuming that the positive charges at NR1 will immediately disappear) (**Fig. 3C, 3D**), electrons 1 and 2 may continue to move toward NR1 due to the residual momentum; for electron 3, it initially will still move toward NR1 but may quickly change its initial direction due to the positive charges of RN2. For electron 4, it will move toward NR2, in a similar manner as electron 1 when NR1 was activated. The movements of electrons 5 and 6 will be similar to the movements of electrons 2 and 3 when NR1 was activated.

Based on the above modeling analysis, it is understood that after the activation (*i*.*e*., AP) disappears at NR1 or NR2, more free electrons will remain inside the segments of a neuro-MT corresponding to the NRs as the positive charges of these small segments will draw free electrons from both directions to these small segments.

### Mechanism of neuro-MT-mediated saltatory neuroelectrical transmission

Based on the above analysis of the EFs and movements of free electrons inside the tunnel of a neuro-MT, the mechanism of neuro-MT-mediated salutatory neuroelectrical transmission is explained below. As schematically depicted in **Fig. 4A**, an axon (which contain bundles of neuro-MTs inside) is often wrapped with a myelin sheath which is interspaced with NRs, and it is known that the APs are initiated at the trigger zone (*i*.*e*., the start region of an axon) and then transmitted in a salutatory manner through the NRs. As explained in the preceding section, prior to the generation of an AP at the trigger zone or NR (*i*.*e*., during the resting state), more free electrons are attracted to the inner surface regions of the neuro-MT segments corresponding to NRs (depicted in **Fig. 4B**). The generation of an AP at the trigger zone is caused by the rapid influx of extracellular Na^+^ through the opening of the fast Na^+^ channels. The rapid increase in cytosolic Na^+^ ions in this small segment of the axon will result in the binding of Na^+^ cations onto a fraction of the negative-charged residues on the outer surface of a neuro-MT, functionally similar to placing a “clamping voltage” on this segment of the neuro-MT. As such, EFs will be generated inside a neuro-MT immediately. First, on the cross sections (*i*.*e*., the ***x–y*** planes), there is an EF with its directions projected away from the central axis (as shown in **Fig. 2C**), which will force the free electrons to leave the inner surface region and move toward the center regions of the tunnel (depicted in **Fig. 4C**).

**Figure 4.**
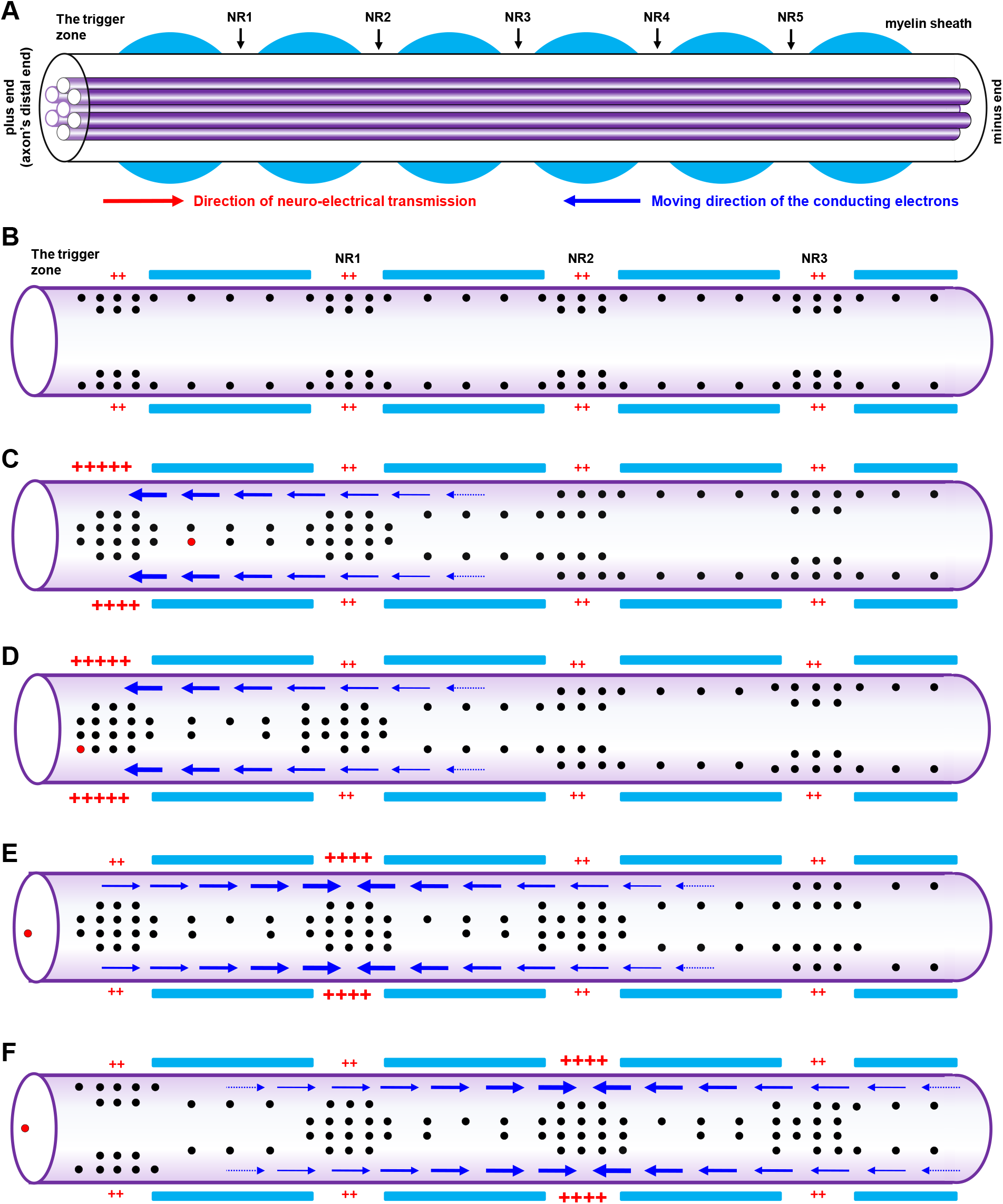
Proposed mechanistic explanation of neuro-MT-mediated saltatory neuroelectrical transmission. **A**. Schematic illustration of an axon that contains bundles of neuro-MTs inside. The left side represents an axon’s distal end, which is close to the neuronal body (soma) and commonly referred to as the “plus end” of a neuro-MT. The right side is the minus end (close to the synapse) of a neuro-MT. The myelin sheath wrapping around the axon (in blue) is usually 1–2 mm in length and separated by a short space (1–5 μm) called “node of Ranvier” (ND). **B**. Distribution of free electrons inside a neuro-MT under the resting condition. The blue colored bars represent the regions where the myelin sheath is located. As depicted, at the resting state, more free electrons gather inside the MT segments corresponding to the NRs, and the free electrons are closely attached to the inner surface region, supposedly in close contact with the positive-charged residues. **C, D**. Movements of free conduction electrons inside a neuro-MT following the generation of an AP at the trigger zone. The formation of an AP in a small segment of a neuro-MT is associated with fast Na^+^ influx, and then the cations will bind to the outer surface of the neuro-MT in that segment, which is functionally similar to placing a ‘clamping voltage’ there. Due to the EF formed on the cross-sections, the free electrons will move toward the center part of the tunnel (as depicted in **C**), and in the meantime, the free electrons on the right (including those at NR1) will begin to move toward the trigger zone (**D**). The migration of some free electrons inside a neuro-MT away from NR1 will result in the release of cations which are initially attracted to the outer surface of the neuro-MT in that segment. The released cations will then increase the free cation concentrations in that part of the cytosol, which will cause a small increase in the resting potential (usually around –70 to –80 mV) inside the nerve terminal and thus trigger an AP (through activation of the voltage-sensitive fast Na^+^ channels in the NR1). Note that the blue arrows indicate the directions of free electron movements, and arrow thickness reflects the relative EF intensity exerted on the free electrons. **0045**. Movements of the free conduction electrons inside a neuro-MT when an AP is triggered at NR1. The clamping voltage at NR1 will draw the free electrons on the right continue to move toward NR1, which will lead to the generation of AP at the subsequent NR2. Meanwhile, the clamping voltage will also exert an opposite EF on the free conducting electrons on the left (they are still moving toward to the trigger zone), and this effect will either slow down their leftward movement or may even change the original direction of movement. As represented by the red electrons, after each neurotransmission, only a very small quantity of free conduction electrons are consumed, *i*.*e*., exiting the neuro-MT and reaching the end pool. **F**. Movements of the free conduction electrons inside a neuro-MT when an AP is triggered at NR2. The sequence of events will be nearly identical to what are seen when an AP is generated at NR1 as described above for **E**.

Second, there is an EF generated along the central axis, which points in the directions away from the positive charge-covered segment (depicted in **Fig. 2B**). This EF will force the free electrons on the right (including those in NR1) to move toward the trigger zone (depicted in **Fig. 3**; also explained in **Fig. 4C**). The migration of free electrons inside a neuro-MT away from NR1 will result in the release of cytosolic cations (such as K^+^) which are attracted to the outer surface residues of that segment during the resting state. The released cations will cause a small increase in the intracellular resting potential (normally around –70–80 mV) in that segment, which will then trigger the generation of an AP in NR1 by activating the voltage-gated fast Na^+^ channels in the NR1 region. Notably, it is known that the plasma membrane at the NRs contain the highest levels of the voltage-gated Na^+^ channels (47-49). The opening of these Na^+^ channels at NRs will be automatically inactivated (*i*.*e*., closed) when the intracellular potential reaches 5–10 mV, and in the meanwhile, the voltage-gated fast K^+^ channels will be activated to efflux K^+^ until the intracellular potential reverts back to the original voltage (49). Afterwards, the Na^+^/K^+^ ATPase is activated to restore the intracellular pools of Na^+^ and K^+^. It is known that an AP only exists for a very short duration and then disappears at each NR, largely due to the ultra-sensitive voltage-gated opening and closing of the fast Na^+^ and K^+^ channels.

Similarly, the AP generated at NR1 will serve as a clamping voltage and will draw the free electrons on the right (*i*.*e*., those at NR2) to move toward NR1 (**Fig. 4E**), which will lead to the activation of the voltage-sensitive Na^+^ channels and generation of an AP at NR2 (**Fig. 4F**). In the meanwhile, the clamping voltage will exert an opposite EF on the conduction electrons on the left of NR1 (which may continue to move toward to the trigger zone), and this opposite force will either slow down their leftward movement or may even make the electrons to move backward toward NR2.

As explained above, when the AP moves from the initial trigger zone to the next NR (*i*.*e*., NR1) in a salutatory manner, the clamping voltage in the preceding NR will quickly disappear and will return to the resting state, which is due to the closing of the inward voltage-gated Na^+^ channels and the opening of the outward voltage-gated K^+^ channels. As a result, the EF on the cross-section of the MT at the preceding NR will return to the initial resting state. Based on the modeling results shown in **Fig. 1D**, the EF on the cross section during the resting state will attract the free electrons toward MT’s inner surface wall. Notably, the alternating positive- and negative-charged residues lining on the inner surface of a neuro-MT (shown in **Suppl. Fig. S3A, 3B, Supple. Fig. S4**) form the spiral dipole ring structures, which serve as effective “speed bumps” and will quickly halt the free electrons from moving in their original directions. After the free electrons come to a complete stop, it is expected that they will be restricted in narrow bands (spiral cycles) very close to the positive-charged residues; as such, free electrons are, in fact, sandwiched between two neighboring spiral bands containing negative-charged residues (**Suppl. Fig. S3B**).

The fact that the movements of free electrons inside the preceding segment(s) of a neuro-MT will be very quickly halted is very different from the conventional electrical transmission inside a metal wire which presumably has evenly-distributed free electrons moving along the entire length of the wire. This unique mechanism will maximize energy saving by reducing the consumption of free conduction electrons during the process of neuroelectrical transmission. In addition, owing to the circular forces exerted nearly evenly on the free electrons moving inside the straight vacuum tunnel of a neuro-MT during neuroelectrical transmission, the conduction of these free electrons will be essentially without significant resistivity, *i*.*e*., they will be moving in a quasi-superconducting manner as proposed earlier (1).

Lastly, it is of note that while all tubulin units in a neuro-MT can produce and release free electrons in a GTP-dependent manner, it is expected that the free electrons are not evenly distributed at the inner surface region of a neuro-MT during the resting state (explained in **Fig. 4**). The modeling results also reveal that the free electrons inside a neuro-MT tunnel will tend to concentrate at the segments corresponding to the NRs, which is consistent with experimental observations that the NRs are axonal segments which have the highest capacitance (50).

## CONCLUSIONS

The present computational modeling work suggests that neuro-MTs, which are distinct structures densely packed inside axons and dendrites, are nanosized biodevices that can mediate physiological neuroelectrical transmission, likely in a quasi-superconducting fashion. The binding of cations (such as the influxed Na^+^ during an AP) onto neuro-MT’s outer surface residues will trigger its transition from a resting state to an active state, and the rapid dissociation of the bound cations will trigger the opposite. The circular dipole ring structures at the inner surface will aid in halting the electron movements almost immediately after the disappearance of an AP at the NRs. Neuro-MT-mediated electrical transmission offers a novel mechanistic explanation for the saltatory conduction of AP along an axon. Lastly, the knowledge gained from this study also sheds light on the rational design of biomimetic superconductive materials in the future.

## DATA AVAILABILITY STATEMENT

All the data of the study are available upon request.

## AUTHOR CONTRIBUTIONS

Conceptualization: B.T.Z.

Research design: Y.X.Y.; B.T.Z.

Methodology: Y.X.Y.

Formal analysis: Y.X.Y., B.T.Z.

Investigation: Y.X.Y., B.T.Z.

Resources: B.T.Z.

Data curation: Y.X.Y.

Writing: Y.X.Y., B.T.Z.

Visualization: Y.X.Y.; B.T.Z.

Supervision: B.T.Z.

Project administration: B.T.Z.

Funding acquisition: B.T.Z.

## SUPPLEMENTARY FIGURES LEGENDS

**Supplementary Figure S1.**
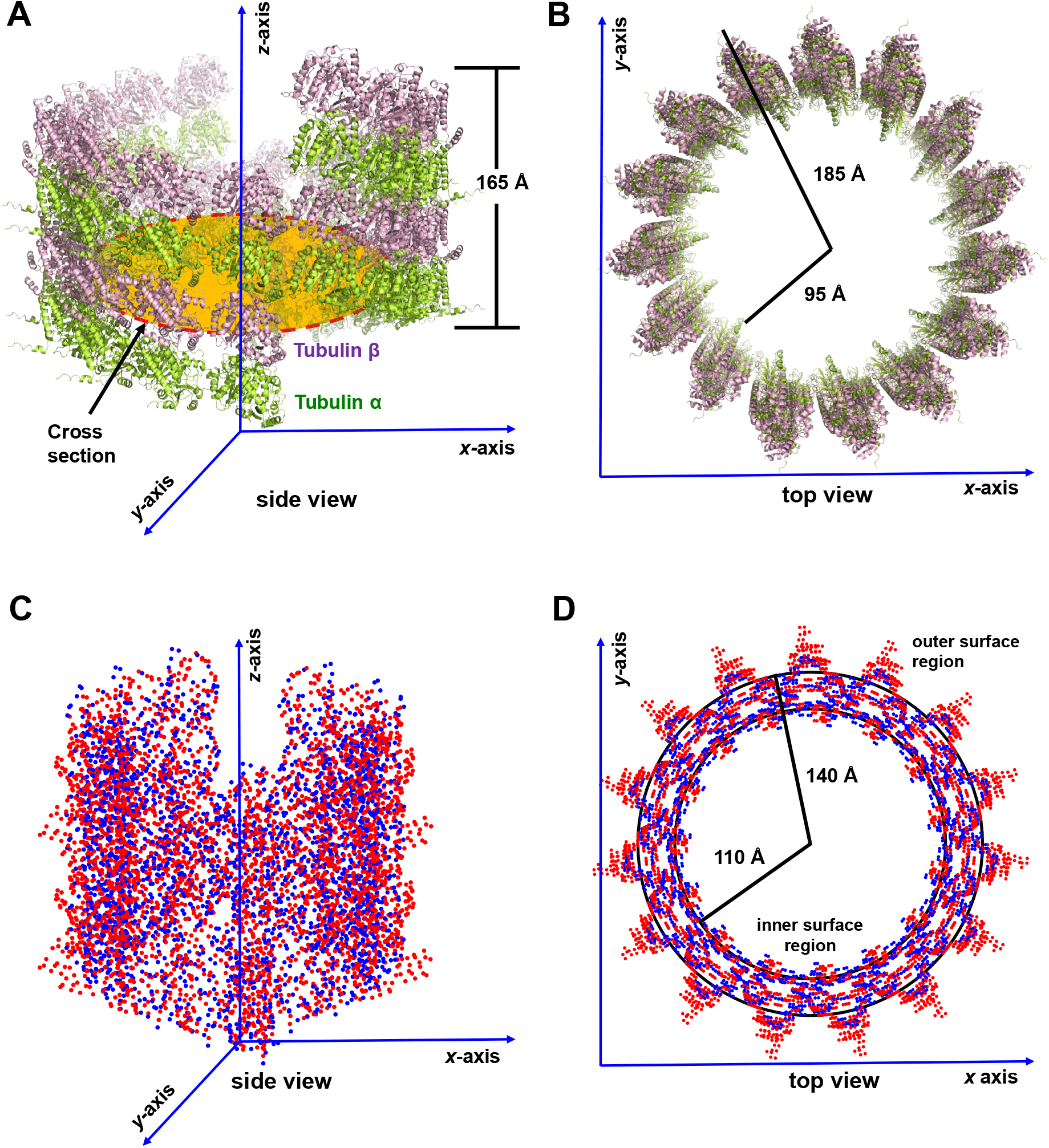
Computationally-modeled structure of a seamless MT and its charge distribution. **A, B**. The side view (**A**) and top view (**B**) of a seamless MT. The tubulin α and β subunits are colored in limon and lightpink, respectively. Note that the structure shown in **A** and **B** is a computationally-modeled MT based the experimentally-determined structure (**PDB code: 6b0i-assembly1.cif; Protein Data Bank**). The missing residues in tubulin α and β are added back according to the AlphaFold-predicted structures. **C, D**. The side view (**C**) and top view (**D**) of the charge distribution of a seamless MT. A red dot represents a negative-charged amino acid residue, a GTP, or a GDP; a blue dot represents a positive-charged amino acid residue or a Mg^2+^.

**Supplementary Figure S2.**
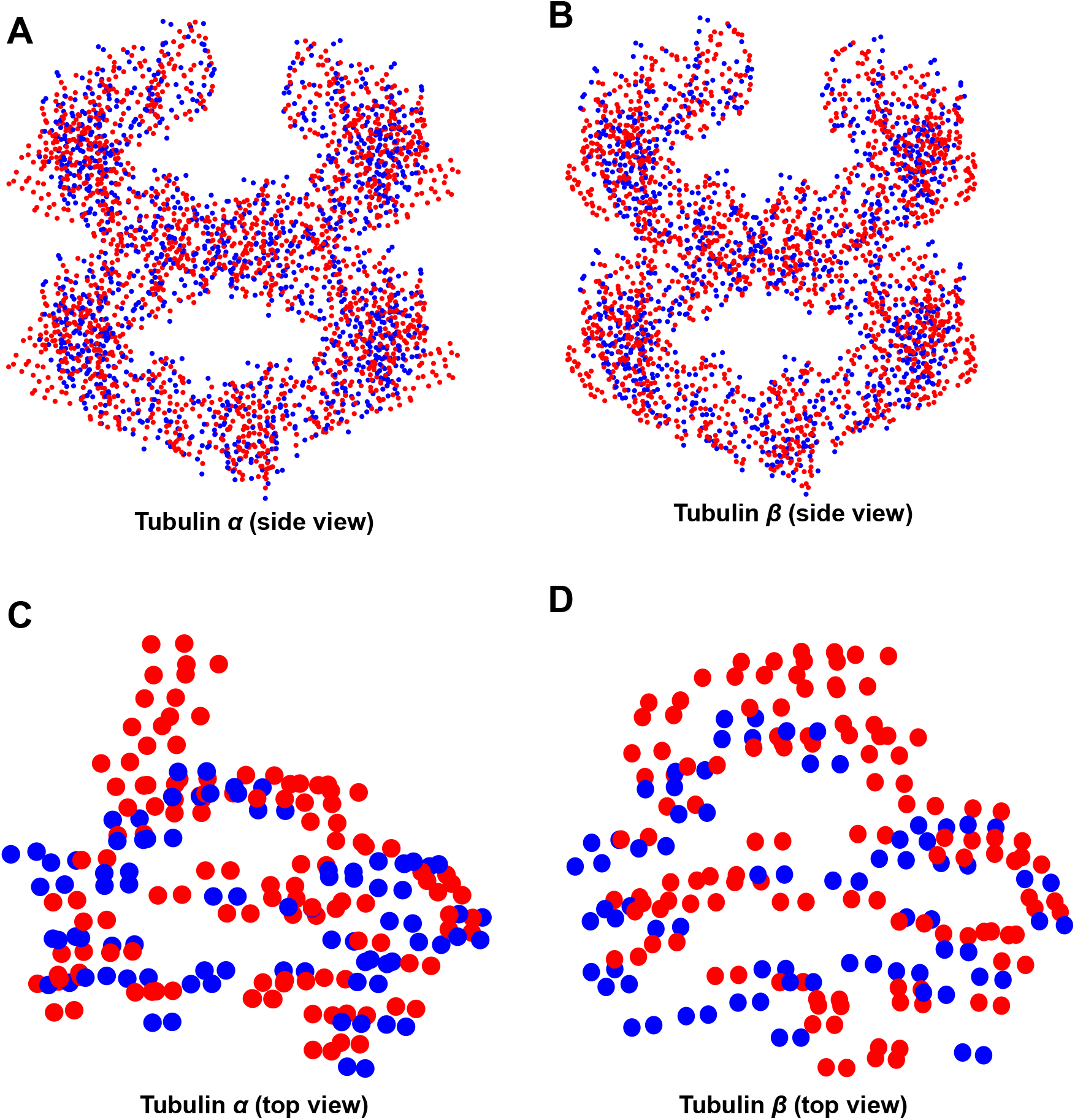
Charge distributions in a 15-membered tubulin α helix (A), a 15-membered tubulin β helix (B), a tubulin α subunit (C), and a tubulin β subunit (D). A red dot represents a negative-charged amino acid residue, a GTP, or a GDP; a blue dot represents a positive-charged amino acid residue or a Mg^2+^.

**Supplementary Figure S3.**
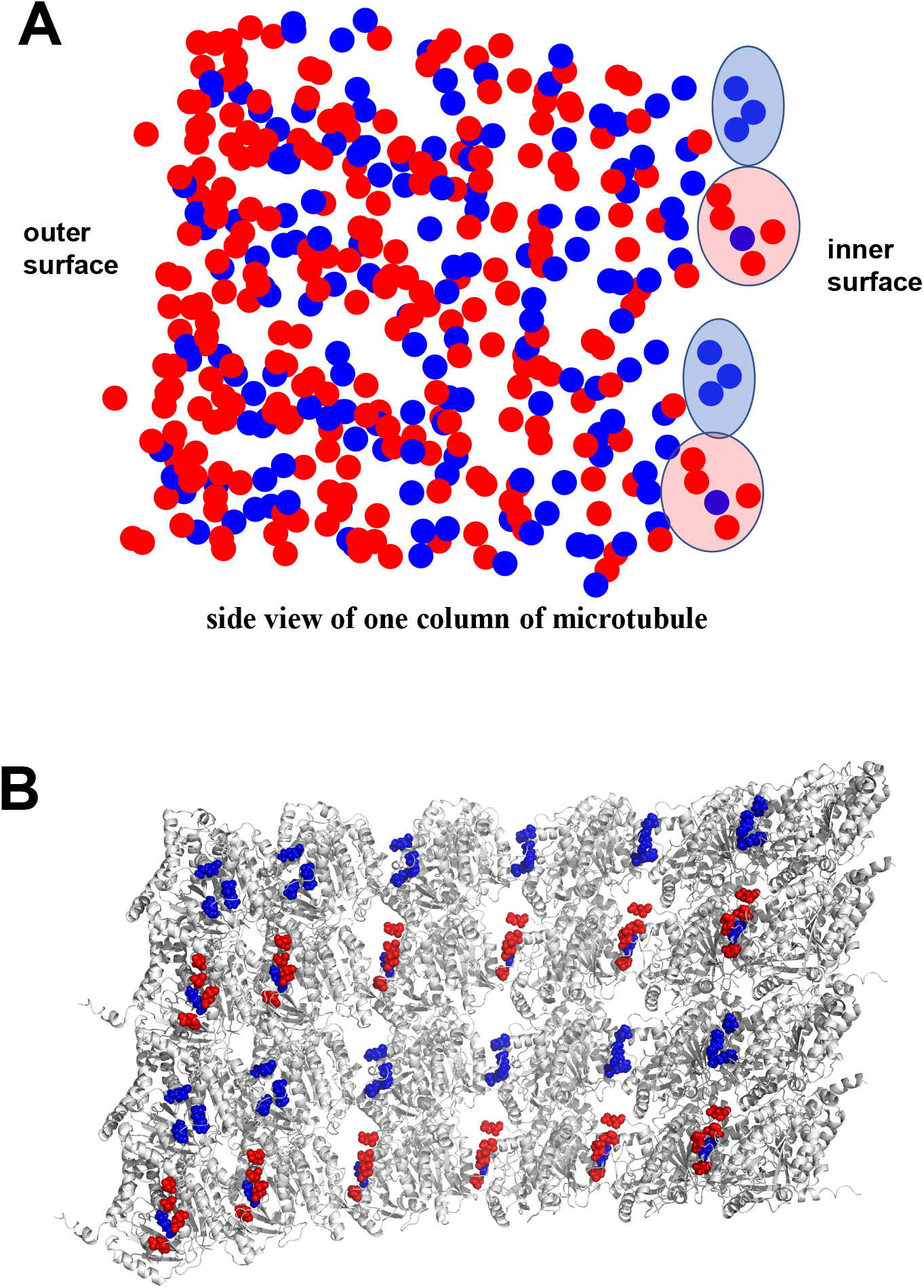
Alternative distribution of negative- and positive-charged amino acid residues on the inner surface of a neuro-MT. **A**. Side view of one column of microtubule. A red dot represents a negative-charged amino acid residue, a GTP, or a GDP; one blue dot represents a positive-charged amino acid residue or a Mg^2+^. **B**. Charged residues on the inner surface in the tubulin αβ dimer structure. The red color for the negative-charged amino acid residues, and the blue color for positive-charged amino acid residues (note that one sphere represents one atom).

**Supplementary Figure S4.**
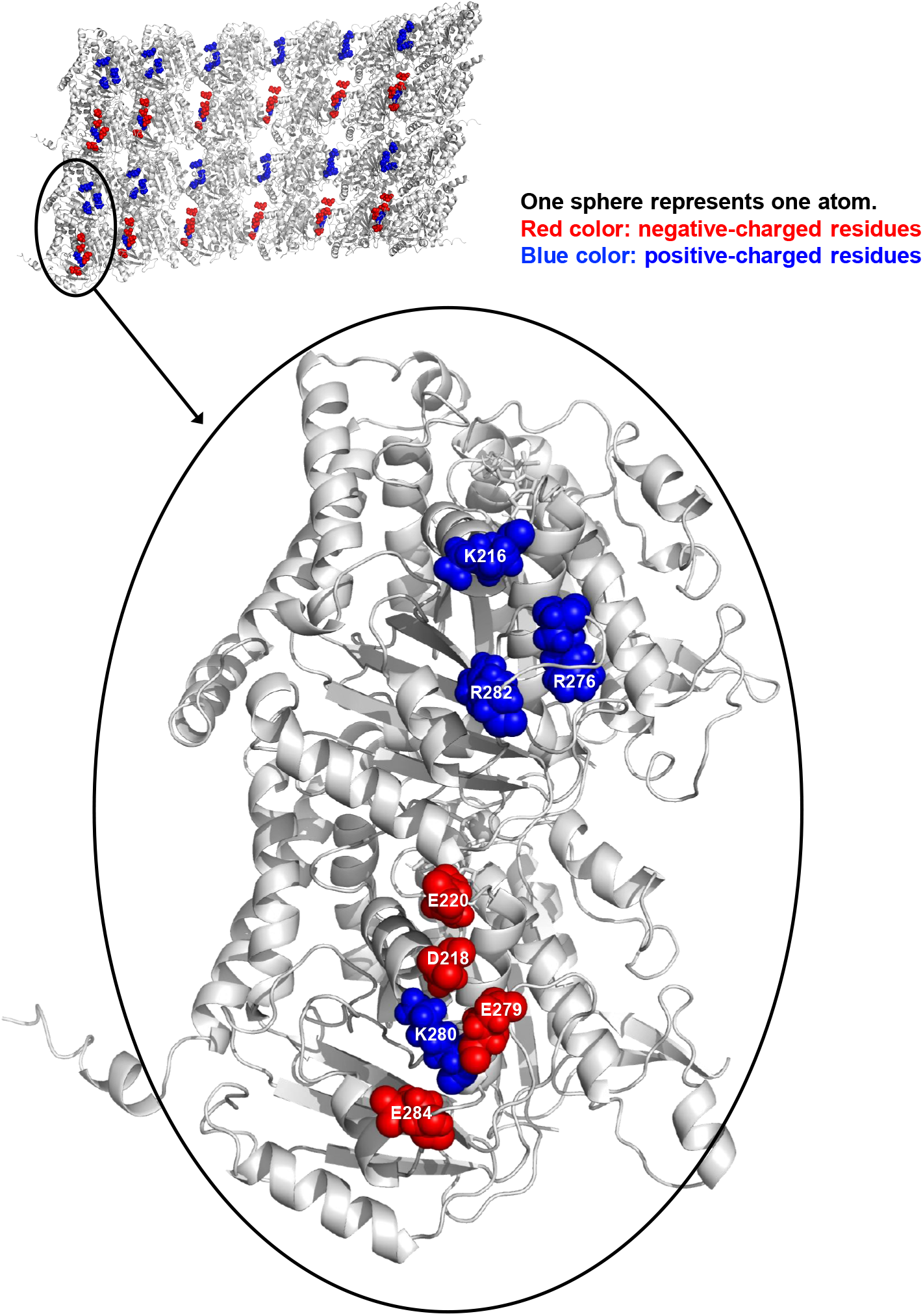
An enlarged view of the alternative distribution of negative and positive-charged amino acid residues on the inner surface of a neuro-MT. The upper left panel shows the charged residues on the inner surface in the tubulin αβ dimer structures (taken from **Supplementary Figure S2B**). The lower panel shows the exact amino acid residues present in the inner surface of tubulin α and β dimer which carry positive (red color) or negative (blue color) charges.

